# Implantable 3D printed multiplexed microtunnels for spinal cord injury treatment

**DOI:** 10.1101/2024.12.02.626000

**Authors:** Sergei Grebenyuk, Volodymyr Medvediev, Oksana Rybachuk, Yevhenii Sheremet, Ibrahim Abdallah, Valeriia Ustymenko, Tetyana Pivneva, Adrian Ranga, Pavel Belan, Nana Voitenko

## Abstract

3D printed scaffolds offer a promising strategy for treating spinal cord injury (SCI). Here we present an innovative biotechnological approach for free-form 3D printing of scaffolds with a biomimetic architecture at a spatial resolution of up to a micrometer, designed for implantation in treatment of SCI in Wistar rats. The fabrication of scaffolds was based on 2-photon photopolymerization of organic polymers and was scalable to lesion geometries. The scaffolds were implemented as multiple densely packed squared parallel microtunnels (50 μm per side) running their entire length. These microtunnels are separated by thin walls (5–10 μm), rendering the scaffolds nearly hollow while maximizing their internal surface area. This design provides an optimal substrate, spatially aligned in the rostro-caudal direction, to support axonal and vascular ingrowth. We have found that the scaffolds, implanted in the excision of the lateral half-fragment of the spinal cord at the low thoracic level demonstrated excellent integration with surrounding tissue without the formation of a significant gliofibrous scar. Myelinated axons and oligodendrocytes, as well as vessels were observed in each microtunnel of the implanted scaffolds in 12 weeks after the operation with at least 1000 axons regenerating in the scaffold throughout its whole length. The treatment significantly improved motor function and reduced spasticity in the ipsilateral paretic limb by 8th week, with recovery sustained for at least 20 weeks. Thus, 3D oriented hollow scaffolds having a large internal surface area and direct continues microtunnels, effectively reducing axonal dispersion, mimic natural structure of the recipient tissue and create conditions for enhancing spinal cord regeneration and recovery of the motor function of the paretic limb.

## Introduction

Spinal cord injury (SCI) is a severe, disabling damage to the nervous system that significantly impairs the quality of life and shortens the life expectancy of the patients (Boakye et al., 2012; Middleton et al., 2012; Geyh et al., 2013). Estimated annual global incidence for SCI is about 1 million cases (James et al., 2019) or by other calculations 10.5 cases per 100,000 persons (Kumar et al., 2018) and estimated global prevalence reached 27 million cases (James et al., 2019). The SCI in most cases accompanied by a significant disability leading to substantial economic losses and creating an additional burden on a state budget (Oliveri et al., 2014). Despite the modest regenerative potential of the central nervous system, it has been demonstrated that spinal cord function can be significantly improved, particularly in cases of severe traumatic injuries. The first priority in treating the SCI is to create optimal conditions for regeneration of nerve fibers across the affected area of the spinal cord. Significant efforts in tissue neuroengineering have been undertaken, involving the technologies of mimicking natural tissue environment by synthetic matrix and supporting axonal regeneration and myelination. For that, a variety of artificially fabricated scaffolds with different chemical compositions have been developed including those based upon peptides, polymers and combinations of synthetic polymers with proteins (Jiu et al., 2024)(Bedir et al., 2020). The scaffolds have been functionalized by different biological active molecules and supplemented with stem cells, including iPSC, primary or genetically modified cells to facilitate the repair of chronic SCI (Li et al., 2019)(Rybachuk et al., 2024).

An increasing body of research highlights the significant impact of scaffold physical characteristics on functional recovery. These features include topology, the ability for axons to extend along the entire scaffold length, material stiffness, adhesion properties to nerve and vascular components, and overall mechanical properties (Mankavi et al., 2023). Ongoing efforts are focused on developing biomimetic microenvironments that can accurately replicate the essential characteristics of the native extracellular matrix in the spinal cord (Jiu et al., 2024).

Despite substantial progress in this field, many challenges remain in achieving effective tissue recovery after SCI. Most suggested matrixes are homogeneous (Kaneko et al., 2015; Palejwala et al., 2016; Chan et al., 2017) (Jiu et al., 2024) and the isotropic nature of the material used for fabrication fails to provide structural guidance for regenerating axons. However, recent advancements in 3D fabrication have enabled the creation of anisotropic scaffolds with one-dimensional orientation. Direct comparison between isotropic and anisotropic scaffolds made from the same material have shown that 3D printing scaffold with an anisotropic structure improve axonal regeneration and promote organized connections within the neural network. This suggests a promising and innovative approach for tissue repair after SCI (Li et al., 2021). Those and other efforts to produce orientated matrixes have resulted in substantial improvements in function recovery after SCI, validating the need for more complex tissue engineering of implants to treat SCI (Nguyen et al., 2017; Wang et al., 2017; Yang et al., 2017; Zhang et al., 2017)(Jiu et al., 2024) (Mankavi et al., 2023).

The one-dimensional oriented scaffolds that provide guidance to axons and vessels can be fabricated using microfibers of varying diameters in a range of 10–100 µm (Zhao et al., 2023), aligned porous materials (Singh et al., 2018) or multiple densely packed parallel microtunnels running the entire length of the scaffold (Koffler et al., 2019). Being tightly packed side by side the microfibers form anisotropic scaffolds that effectively guide axons and blood vessels during spinal cord repair, significantly improving motor function of rats after SCI (Zhao et al., 2023). However, geometrical morphological considerations indicate that tightly packed cylindrical fibers occupy approximately 80% of the cross-sectional area of the scaffolds, leaving only about 20% free space, which can impede tissue recovery after SCI. In the porous materials, the pores may not be well interconnected, and even when they are, anisotropic pores might not create continues and direct pathways throughout the entire length of the scaffolds, which is unfavorable for nerve regeneration (Taylor and Haycock, 2022)(Hibbitts et al., 2022).

Thus, it seems that the microtunnel scaffolds being intrinsically continuous and nearly hollow may provide better guidance for axonal and vascular ingrowth compared to the fibers and pores of anisotropic materials. It is obvious that the scaffold material should occupy as little volume as possible leaving more space for the recipient tissue. However, walls of tunnels, fibers, and fillers, within the scaffolds, limit the space necessary for growing axons and blood vessels. In most fabricated scaffolds the scaffold materials occupy more than 50 % of the inner space (Jiu et al., 2024). Therefore, it is advantageous to design scaffolds with thin yet durable walls that have the smallest possible cross-sectional area. Three-dimensional scaffolds having small sized tunnels and thin walls would also provide a large internal surface area, offering physical support and a spatially oriented substrate for the axonal and cellular growth necessary for successful spinal cord repair. Additionally, smaller tunnel sizes could effectively reduce axonal dispersion, promoting more directed and organized axonal growth that mimics natural structure of the recipient tissue. Small diameter tunnels, with dimensions comparable to those of spinal tracts, could partially replicate the unique morphology of the spinal cord, thereby enhancing neurogenesis and nerve regeneration. However, one-dimensional oriented scaffolds with tunnels smaller than 200 µm and thin walls have never been employed for spinal cord repair, mainly due to technological challenges related to their fabrication (Jiu et al., 2024).

Scaffolds with thin walls of oriented tunnels having a high surface-area-to-volume ratio and being nearly hollow should be manufactured from durable materials to ensure physical stability during and after implantation. Polyethylene glycol (PEG) possesses high mechanical strength, which can be further enhanced by combining it with the polyacrylate crosslinking agent pentaerythritol tetraacrylate (PETA) to create mechanically stable 3D structures. This combination not only improves mechanical stability compared to pure PEG but also enhances cell attachment and proliferation on the surface of printed scaffolds (Klein et al., 2011). For these reasons, this material combination was utilized in the present study.

We hypothesized that 3D printed scaffolds with aligned, continues microtunnels having a size of 50 μm and thin walls between them would be beneficial for axonal and vascular growth, leading to effective regeneration after SCI. To test this hypothesis, we fabricated the scaffolds using high-resolution two-photon photoprinting. We suggested that these 3D-printed scaffolds would mimic the microarchitecture of the spinal cord and provide for a large surface area essential for supporting axonal and vascular regrowth. Immunohistochemical evaluations of the implanted scaffolds revealed considerable axonal, cellular and vascular regeneration after SCI while the behavioural testing demonstrated significant functional motor recovery and reduced spasticity. To the best of our knowledge, this is the first study, in which nearly hollow, oriented multitunnel scaffolds with a high surface-area-to-volume ratio were 3D-printed with a micron resolution and evaluated *in vivo* for recovery after SCI. Additionally, this is the first report to directly demonstrate the critical importance of the physical microstructure of 3D scaffolds for spinal cord regeneration.

## Methods

### Design and fabrication of implants

The scaffolds were designed in a computer aided design software (Solid Edge ST9, Siemens) as a multitude of parallel microtunnels of squared cross-section (50 μm wide) with thin (5 μm) walls between the microtunnels “packed” within a cylinder or semicylinder (Fig. 1). The scaffolds were micro-fabricated using high-resolution 3D printer (Photonic Professional GT2, Nanoscribe GmbH) equipped with 2-photon femtosecond laser. CAD model of the scaffold was pre-processed by DeScribe software (Nanoscribe GmbH) to produce a printing process file with defined printing parameters. A custom-formulation of photopolymerizable resin was used to print implants which contained 2% 2-Benzyl-2-(dimethylamino)-4’-morpholinobutyrophenone (Irgacure 369), 90% poly(ethylene glycol) diacrylate, MW700 (PEGDA 700) and 10% pentaerythritol triacrylate (PETA). Irgacure 369 was purchased from Tokyo Chemical Industry Ltd, the other components were from Sigma-Aldrich. After fabrication, implants were washed 3–4 days in 2-propanol and then kept in ethanol until use. The solvents were refreshed daily for the first week and every week afterwards.

**Figure 1.**
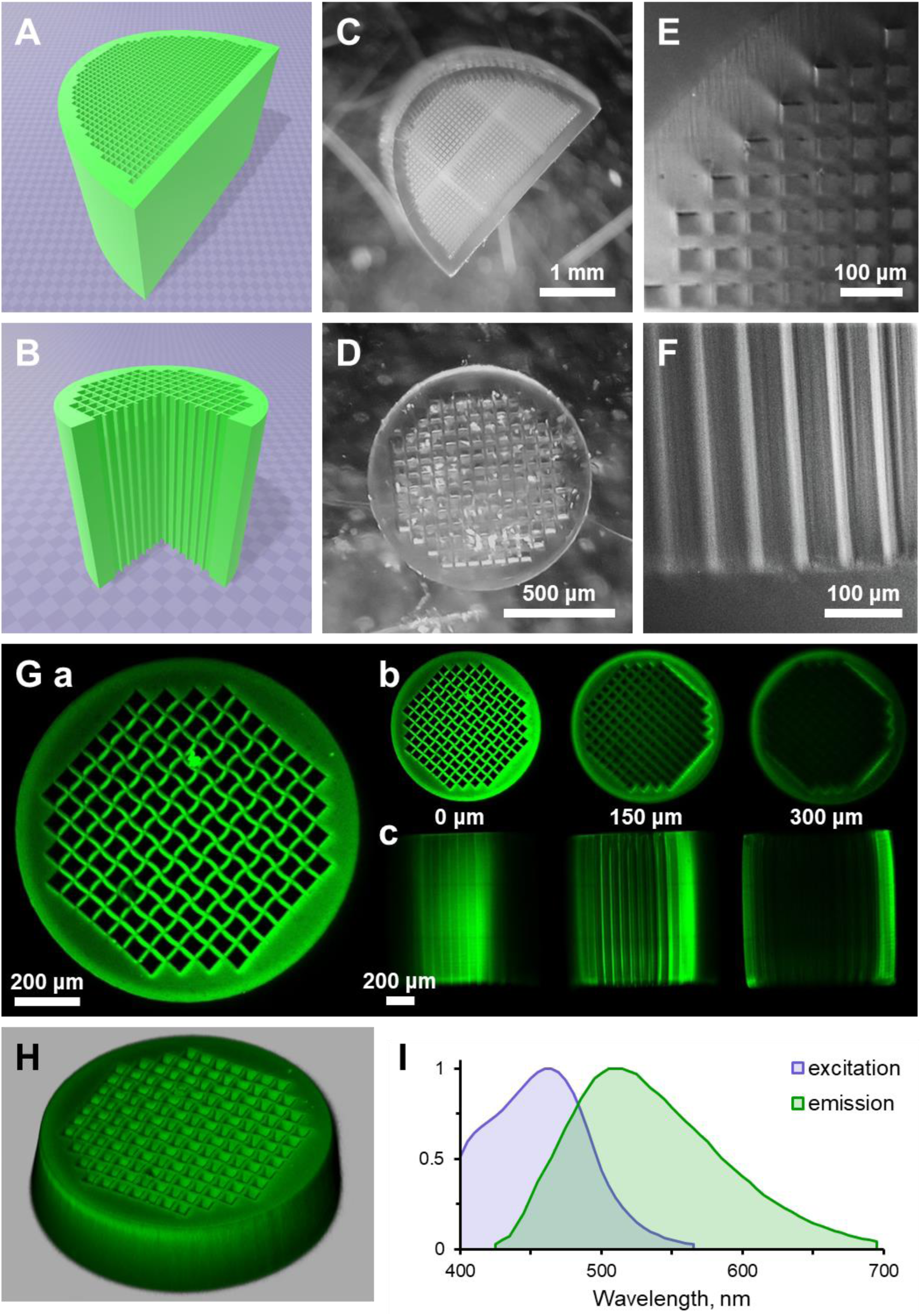
Fabrication and characterization of 3D printed scaffolds with continues oriented microtunnels and thin walls. **A**, **B**. 3D designs for nearly hollow semicylindrical and cylindrical scaffolds used for implantations in a rat spinal cord after hemidescection. **C**, **D**. Images of the scaffolds fabricated by a high-resolution 2-photon 3D printer. Images obtained using an Olympus SZ-51 stereomicroscope. **E**, **F**. Higher magnification axial (**E**) and side (**F**) images of the scaffolds obtained using infrared oblique illumination (Safronov et al., 2007). The most superficial row of microtunnels is shown in **F**. **G**. Fluorescent images of 3D printed cylindrical scaffold obtained using confocal microscopy (Olympus FV1000). The combination of materials used for the photoprinting, PEGDA/PETA, revealed strong green fluorescence when excited in UV range. **Ga**. An axial Z-stack of the scaffold. Z-stack images obtained at 0, 150 and 300 μm depth in axial (**Gb**) and side (**Gc**) projections of the scaffold. Note impossibility to image the scaffold deeper than 150 µm due to light scattering in the scaffold material. Side images of the scaffold (**Gc**) demonstrating high quality of printed tunnels and their continuity throughout the whole length of the scaffold. **H**. 3D Z-stack reconstruction of the first 200 µm of the scaffold. **I**. Fluorescence spectra of scaffold material. Note that the scaffold material demonstrated the almost negligible fluorescence when excited above 543 nm.

### Surgical procedures

Ethics statement. All experimental procedures were approved by the Animal Ethics Committee of Bogomoletz Institute of Physiology (Kyiv, Ukraine) and performed in accordance with the EU Directive 2010/63/EU for animal experiments, ethical guidelines of the International Association for the Study of Pain, and the Society for Neuroscience Policies on the Use of Animals and Humans in Neuroscience Research.

The study was performed on 19 male Wistar rats, aged about 1 month and weighing about 50 g at the time of implantation. The animals were kept in the usual physical conditions, at a constant temperature of 18–20°C under a natural circadian light cycle and fed with a balanced combined feed *ad libitum*. The animals were divided in three groups: 1) an excision group, in which hemi-excision of a left lateral half fragment of the spinal cord having a length of 1 mm in a rostrocaudal direction was performed in the lower thoracic region (n = 8); 2) a group with cylindrical scaffolds, in which the hemi-excision followed by an immediate filling of the lesion site with a cylindrical (1 mm in diameter and 1mm length) 3D printed biomimetic microtunneled scaffold fabricated from PEGDA/PETA (n = 8) and 3) a group with semicylindrical scaffolds, in which the hemi-excision followed by an immediate filling of the lesion site with a semicylindrical (3 mm in diameter and 1mm length) 3D printed biomimetic microtunneled scaffold fabricated from PEGDA/PETA (n = 3).

The hemi-excision of the spinal cord at the T12–T13 level was performed as previously described (Medvediev et al., 2021). Briefly, rats were deeply anesthetized with ketamine and xylazine (70 mg/kg and 15 mg/kg, respectively, intraperitoneal injection), confirmed with the lack of the corneal reflex and loss of hindlimb pain sensitivity. A longitudinal skin incision was made within Th8–L2 vertebrae and muscles were retracted to expose the spinal column. Left-lateralized laminectomy was performed at the level of T11 or T12 vertebrae. After spontaneous stopping of epidural venous bleeding, the spinal cord was punctured as close as possible to the left edge of the posterior median vessels through the ventral direction with the help of an insulin syringe needle maintaining a perpendicular direction of the needle to the dorsal surface of the spinal cord. In a similar way, through punctures were performed ∼0.5 mm more rostral and more caudal. One branch of an open ophthalmic scissors was inserted into every two adjacent punctures, and a longitudinal paramedian spinal cord dissection was made at a distance of ∼1 mm in several steps. One of the branches of the ophthalmic scissors was successively inserted into the rostral and caudal end of the longitudinal wound and the left half of the spinal cord together with the nerve roots was covered with the second branch and transacted in several steps. A fragment of the left half of the spinal cord cut out in this way from three sides with a length of ∼1 mm was removed with ophthalmic tweezers forming a lesion site. In the animals of groups 2 and 3 the respective 3D printed scaffolds were implanted into the lesion site. Immediately before implantation, the scaffold was transferred to an isotonic solution of sodium chloride, then to a freshly prepared isotonic solution of sodium chloride and 0.1 % hydroxymethylquinoxaline dioxide (Dioxydine, PJSC Farmak, Ukraine) for ∼40 min. The excess, dropwise part of the antibacterial solution around the matrix was removed by touching the muscle or fascia within the surgical wound.

In animals of all experimental groups, the access window to the spinal canal was covered with a fragment of subcutaneous fascia. After that, soft tissues and skin were sewn by means of two rows of knotted sutures. The wound site was treated with a povidone-iodine solution (EGIS, Hungary). For the prevention of infectious complications, a solution of bicillin-5 (JSC “Kyivmedpreparat”, Ukraine) was injected into the posterior cervical region subcutaneously at a dose of 0.5 million IU/kg. Intraperitoneal administration of a solution of dexamethasone (“KRKA”, Slovenia) at a dose of 5 mg/kg was used for anti-inflammatory and antiedematous therapy. After surgery, until complete awakening, the animals were kept at increased air temperature. Subsequently, the animals were kept in individual deep plastic cages at an average air temperature of 18–20°C.

### Immunohistochemistry

Spinal cords were harvested in three (1 rat) and five (5 rats) months after implantation. For that, animals were deeply anesthetized with diethyl ether and perfused intracardially with cold (4° C) solution of 4% paraformaldehyde in 0.1 M phosphate buffered saline (pH 7.4). Solid vertebral columns (Th9–L4) were removed and postfixed in 4% paraformaldehyde in phosphate buffered saline at 4° C during one week. After that, spinal cords were removed from the vertebral canal. Spinal cords were sectioned longitudinally into 70–100 μm slices using Campden 752M vibrotome (Campden Instruments, UK). The slices were incubated in blocking solution containing 0.5% bovine serum albumin (BSA) and 0.3% Triton X-100 in 0.1 M phosphate buffer (PB) for 2 hours at room temperature. Immunohistochemical staining of slices was carried out to visualize axons (cytoskeleton protein β-tubulin) and myelin sheaths (Myelin Basic Protein, MBP). For that, mouse monoclonal Anti-β-Tubulin Isotype III antibody (Sigma Aldrich, T5076) and rabbit polyclonal antibody to MBP (Novus Biological, UK, NB110-79873SS) were diluted 1:250 in PB with 0.5% BSA and 0.3% Triton X-100. Sections were incubated with these primary antibodies for 24 hours at 4°C followed by incubation with secondary antibodies, including Alexa Fluor 555 donkey anti-mouse (Invitrogen, A31570, at 1:1000) and Alexa Fluor 647 goat anti-rabbit (Thermo Fisher Scientific, A27040, at 1:1000). Secondary antibodies were incubated for at least 2 hours at room temperature. Some samples were counterstained for 10 min at RT with Hoechst (Sigma, USA, at 1:5000) to visualize cell nuclei. After three washes, sections were mounted with VECTASHIELD mounting medium (Vector Laboratories, Burlingame, CA) and inspected using a confocal microscope (FV1000-BX61WI, Olympus, Japan). One to three slices per animal were quantified. Quantification was performed using FV10-ASW software (Olympus, Japan). Excitatation and emission spectra of scaffold material were measured using Till Photonics imaging system (Till Photonics, Germany).

### Behavior testing

*Functional motor test*. The Basso–Beattie–Bresnahan (BBB) scale (Basso et al., 1995) with some modifications was used for the assessment of motor function of hindlimbs in each animal. Our scoring was based on the BBB rating scale for the incomplete spinal cord injury in rats keeping in mind technical limitations of the BBB-scale for the evaluation of unilateral functional deficit (Medvediev et al., 2021). Tested animal was allowed to freely move and was evaluated for its hindlimb joint movements, paw placement, weight support, and limb coordination according to the scoring criteria.

The BBB open-field 21-point locomotion rating scale was assessed weekly during first 8 weeks and each 4 weeks after that over a total 20-week period after the implantation. It was done by the observer blinded to group identity. The BBB scores for both hindlimbs was determined during the observation of motor activity on a horizontal open solid surface sufficient for the study of the long-term unidirectional step locomotion. The calculation of functional scores was carried out no earlier than on the 6^th^ day after the injury in view of the ethical regulations for working with experimental animals. The value of functional scores immediately after modeling the injury was conditionally assumed to be zero (state of spinal shock). When signs of both adjacent values of BBB score were detected during testing, the average, half value was recorded.

*Assessment of spasticity in a paretic limb.* The spasticity score at the level of hip (manual abduction of the limb to the side), knee (manual extension during abduction of the limb to the side) and ankle joints of the posterior ipsilateral limb was evaluated according to the Ashworth scale in the interpretation of Dong et al. (Dong et al., 2005)(Kopach et al., 2017) in our technical modification (Medvediev et al., 2021), recording the maximum value among the three manually studied joints of the hindlimb. Assessment of spasticity was carried out no earlier than on the 6^th^ day for ethical reasons. The value of Ashworth scale immediately after modeling the injury was conditionally assumed to be zero (state of spinal shock). When signs of both adjacent values of Ashworth scale were detected during testing, the average, half value was recorded.

The results of functional testing were presented according to a conditional time scale.

### Statistics

Two-group comparisons of immunochemical data were tested by two-tailed Student’s t-test using Origin software (OriginLab Corporation, United States). Multiple-group comparisons were tested by one way ANOVA, followed by *post hoc* analysis (if necessary) using Student’s t-test (OriginLab Corporation, United States). The Kolmogorov–Smirnov test was used to estimate the normality of the data.

Statistical processing of data for monitoring BBB and Ashworth scores of the paretic limb was carried out using STATISTICA 10.0 software (TIBCO Software, United States) and R (The R Foundation). The non-parametric one-tailed Mann–Whitney U test was used for the comparative evaluation of the scores between the groups. The reliability of the difference in BBB and Ashworth scores at different time points within a group was assessed by Wilcoxon matched pairs test. ANOVA or Student’s t-test were used for comparisons of mean values of the groups if sample data were normally distributed. Shapiro–Wilk test was used to check if the normality assumption can be applied to the data. The averaged values in this case were presented in the form of mean ± S.E.M. If other is not indicated, the assumption regarding the statistical significance of the obtained result was considered true if the probability of the null hypothesis was less than 0.05 (p < 0.05).

## Results

### Fabrication and characterization of 3D printed scaffolds with continues oriented microtunnels and thin walls

High-resolution 2-photon printing allows the scaffold fabrication of arbitrary design that may fit the anatomy and microarchitecture of a particular SCI. In this work, the spinal cord lateral hemi-excision at the lower thoracic level was used as a SCI model (Medvediev et al., 2021). This model involves the unilateral hemi-excision of 1 mm of the spinal cord and disrupts all ascending and descending tracts on one side only. The hemi-excision produces a highly selective and reproducible SCI, which is convenient and useful for studies of spinal cord regeneration and functional recovery. We hypothesized that scaffolds having a large internal surface area and small volume of material designed as a set of continues microtunnels with thin walls between them are essential to retain physical support across a lesion site and to align new vessels and regenerating axons across the lesion.

We therefore designed scaffolds having different shapes and a length of 1 mm as a regularly spaced network of microtunnels whose thin walls were made of a hydrogel allowing free diffusion of dissolved molecules. (Fig. 1). 3D prints with the 2-photon Nanoscribe printer were performed as described previously (Grebenyuk et al., 2023). We used a custom formulated photo-polymer based on a mixture of polyethylene glycol diacrylate (PEGDA) and photocrosslinker pentaerythritol triacrylate (PETA) having sufficient porosity and cell-adhesion as well as enabling rapid diffusion (Grebenyuk et al., 2023). 90% of PEGDA and 10% of PETA were used in the polymer since it has been shown that this proportion is optimal for the protein-adsorbing and cell-binding characteristics of respective hydrogel (Klein et al., 2011). The hydrogel permeability to water-soluble molecules was previously estimated using diffusion of fluorescent molecules. It was shown that the three-dimensional diffusion of fluorescein inside matrices with a characteristic size of 2 mm made of the hydrogel was observed in less than 10 min (Grebenyuk et al., 2023). However, tissues regenerating within the scaffolds requires metabolic support not only from outside of the scaffold but also and mainly from capillaries, which could be ingrown in the microtunnels. The average intercapillary distance in the CNS is about 40–50 μm (Duvernoy et al., 1983)(Nicholson, 2001) and, therefore, we decided to print the microtunnels of similar sizes suggesting that if one capillary were ingrown in one microtunnel it would provide for the necessary metabolic support, A small distance between walls of the microtunnels (40–50 μm) would simultaneously result in a large internal surface area necessary for physical support across a lesion site. The combination of 2-photon printing with the PEGDA/PETA hydrogel allowed printing of a variety of scaffolds including cylinders (a diameter of 1.0 mm and a length of 1.0 mm) and semicylinders (a diameter of 3.0 mm and a length of 1.0 mm) (Fig. 1A–D) with different shapes of microtunnels with their wall thickness from 5 to 10 µm (Fig. 1E, F). The scaffolds with squared microtunnels with a side of 50 μm and wall thickness of 5 μm were used for implantations in sites of SCI (Fig. 1G, H). This design of scaffolds produces about 10 times larger surface area to support regeneration per unit of cross-section area compared to one utilizing round tunnels with a diameter of 200 μm (Koffler et al., 2019).

Quality of printing and fluorescent properties of scaffolds were tested using the confocal microscopy. Excitation of scaffolds photofabricated from PEGDA/PETA using a UV laser at 351 nm resulted in strong green fluorescence allowing to visualize the scaffold microstructure (Fig. 1G, H). The end view of the very top part of the scaffold demonstrates the high quality of printing using 2P technology. A stack of the images was acquired from the top up to 300 μm deep inside the scaffold. The representative images from the stack and 3D reconstruction of the end view of the scaffold are shown in Fig. 1G, H). The data clearly demonstrate that microtunnel structure is well-preserved up to 150 μm deep from the top (Fig. 1Gb). At the same time, the strong scattering of excitation laser beam did not allow the light to penetrate deeper that prevents checking of microtunnel continuity throughout the whole length of the scaffold. Besides, it appears that the high level of fluorescent intensity of scaffold material will interfere with a possibility to use fluorescent tags of secondary antibodies, which are excited in the UV range of wavelengths. In order to check the continuity of the microtunnels, the side view images of the scaffolds were obtained (Fig. 1Gc). The continuity of the microtunnels and high quality of printing are clearly visible in at least the upper 150 μm of the scaffolds. The same results were obtained when different side views of the same scaffolds were systematically investigated (data not shown). Thus, the quality control of 3D printing showed that in the most part of the scaffold volume (150 μm deep from the ends and side of the cylinder of 1 mm in diameter and 1 mm in length) the microtunnels are continuous throughout the whole scaffold length and had no visible defects at the micron level of resolution. In order to check the quality of printing in the central part of scaffolds, it was attempted to cut the scaffolds in slices using a vibrotome. Unfortunately, the obtained slices with a thickness from 50 to 200 μm were partially damaged by the employed slicing procedures that did not allow to validate the printing quality in the central part of the scaffolds.

After implantation of scaffold in sites of SCI, the certain ingrowth of recipient tissue including vessels and nerve fibers is suggested based on the results of earlier studies (Rybachuk et al., 2024). After sacrificing animals, the spinal cord together with the implanted scaffolds were removed and subjected to immunochemical investigation. The fluorescence of the scaffold material (Fig. 1G, H) creates a good opportunity to visualize the scaffolds but simultaneously may prevent observing the immunochemically-stained tissues inside the scaffolds. Therefore, we studied the excitation and emission spectra of the scaffold material in order to properly chose fluorescent dyes for the secondary antibodies for the following immunochemistry. The emission spectrum of the scaffold material when excited at 405 nm shown in Fig. 1I demonstrates that it propagates up to 700 nm obstructing simultaneous confocal imaging using longer wavelength lasers if UV excitation is engaged for scaffold visualization. However, in case of sequential excitation of fluorophores it still can be possible to separately visualize both the hydrogel matrix and recipient tissue if the scaffold material is not excited at longer wavelengths. The excitation and emission spectra of printed scaffolds were measured using a spectrofluorimeter demonstrating low excitation above 500 nm (Fig. 1I). Additionally, using the confocal system, it was found that the emission was negligible at 633 nm excitation even at 100% of the laser power; the low level of emission was observed at 543 nm excitation when high laser power above 30% was engaged. At the same time, the scaffold material was strongly fluorescent when excited at 405 nm and 488 nm (Fig. 1I). Thus, in order to observe the scaffold material and recipient tissues inside the scaffolds, the dyes excited at 543 and 633 nm were employed while to examine the scaffold structures 351, 405, 488 nm lasers were used.

### Scaffold implantation and removal

Scaffolds were implanted into rats right after the hemi-excision of the spinal cord. In order to produce the hemi-excision, Th8–L2 segments and muscles were retracted to expose the spinal column. Left-lateralized laminectomy was performed at the level of segments T12–T13 cutting the spinal cord out from three sides to form a defect of about 1 mm sufficient to keep the scaffold tightly fixed by its ends (Fig. 2A, B). Eight Wistar rats underwent placement of 3D cylindrical biomimetic microtunneled scaffolds (a diameter of 1.0 mm and a length of 1.0 mm) into the lesion site (Fig. 2D); three received (1.5 × 3.0 x 1.0 mm) 3D semicylindrical microtunneled biomimetic scaffolds (a diameter of 3.0 mm and a length of 1.0 mm) (Fig. 1A–D); eight controls underwent the hemi-excision only (Fig. 2C).

**Figure 2.**
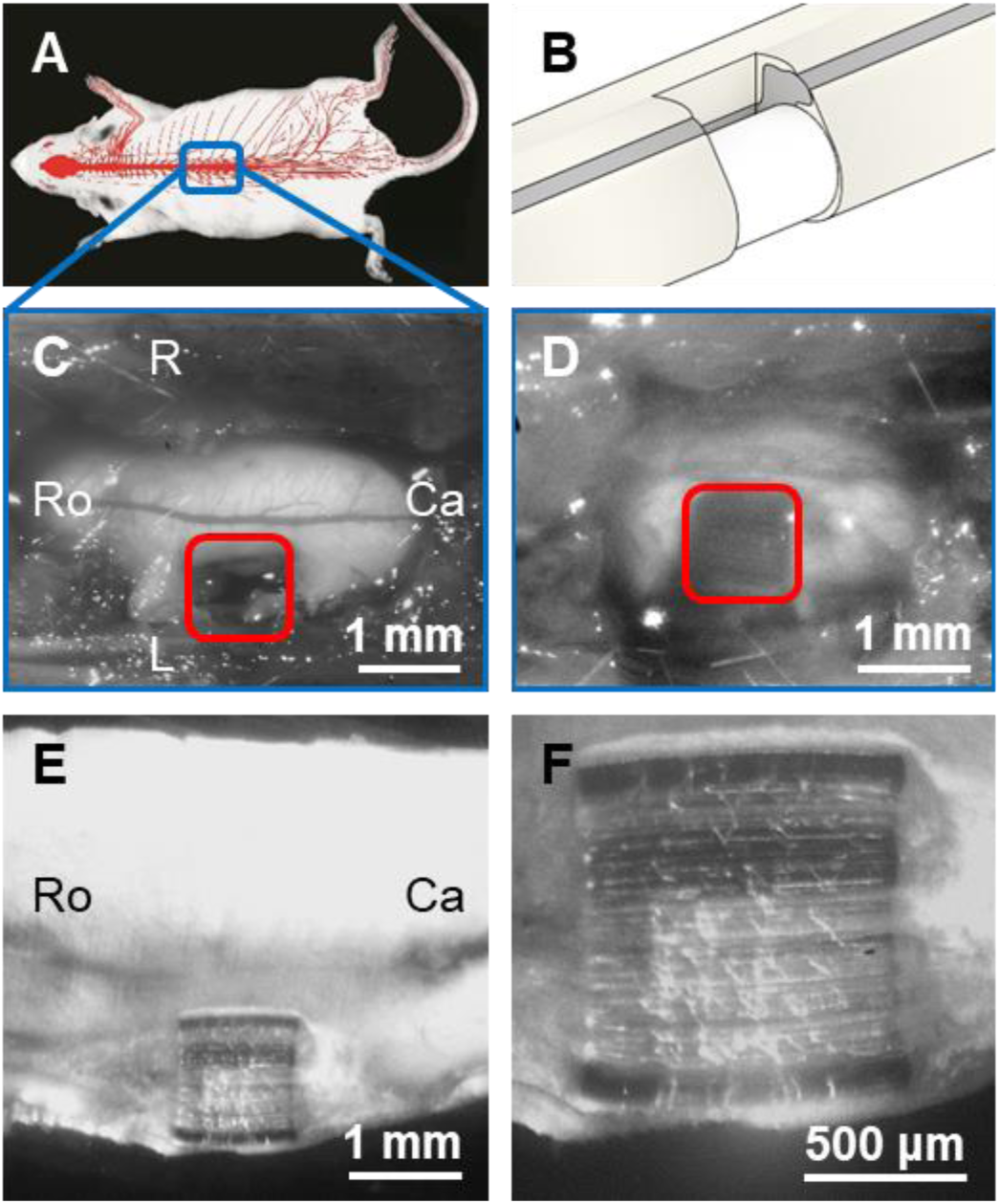
Scaffold implantation and removal. **A**. A picture demonstrating a site at the T12–T13 level where the hemi-excisions of the spinal cord and implantations of the scaffolds were performed (modified from http://larrywswanson.com/?page_id=1148). **B**. Drawing showing the position of the cylindrical scaffold at the implantation site in the spinal cord. **С**. The spinal cord with a left side lesion site of 1 mm in rostocaudal direction. **D**. A cylindrical scaffold (1 mm in diameter and 1 mm in length) implanted in the lesion site. Red boxes in **C** and **D** indicate the sites of hemi-excision. **E**. A longitudinal slice of spinal cord with the implanted scaffold. The spinal cord was harvested for slicing and following immunochemistry in 12 weeks after scaffold implantation. **F**. Higher magnification of the same slice demonstrating integration of scaffold in the recipient tissue, retaining its structure and some damage of microtunnels produced by a cutting procedure.

Initially we suggested that the semicylindrical scaffolds covering the entire cross-section of the spinal cord would lead to the better regeneration of axons and better functional recovery compared to the cylindrical ones. However, it turned out that the semicylindrical scaffolds in all tested animals (n = 3) could not be properly fixed inside the spinal cord hemi-excision and completely or substantially protruded from the lesion site during the time course of operation. Taking this result into account we did not conduct further implantations of the semicylindrical scaffolds.

All control animals and ones underwent placement of cylindrical scaffolds survived the first experimental period of 12 weeks after implantation, and no sign of infection was observed. During this period the animals were monitored each week for functional motor recovery and spasticity.

One rat implanted with the scaffold was sacrificed after 12 weeks and used for immunohistochemical study. During the second experimental period (weeks 13–20) the animals were tested each four weeks. After 20 weeks of surgery, the remaining animals were sacrificed and scaffolds with flanking recipient tissue of the spinal cord were harvested and immunohistochemically analyzed for nerve and vessel regeneration. After both 12 and 20 weeks *in vivo*, 3D biomimetic scaffolds retained their initial structure. The harvested spinal cords with the embedded scaffolds were cut in longitudinal slices with a thickness of 70–100 μm consisting both ipsi- and contralateral parts of the spinal cord (Fig. 2E). The scaffolds were well aligned along and integrated within the ipsilateral part of spinal cord (Fig. 2E, F). The site of hemi-excision was completely occupied by the scaffold and no lesions were observed around while the hemi-excision alone resulted in the permanent severe lesion spreading 1–3 mm to the rostral and caudal directions in 20 weeks after the injury (data not shown). The scaffolds were well preserved without breakage or deformation retaining physical support across a lesion site during the whole long period of implantation. However, the cutting procedure necessary to study the tissue regeneration inside the scaffolds damaged the microtunnels as evident by irregular wrinkles observed over the surface of sliced matrix (Fig. 2E, F).

### Morphological assessment of tissue ingrowth in the scaffolds implanted in a site of SCI

The cytoskeleton of the axons contains straight parallel bundles of extended but discontinuous microtubules running all along consisting of polar polymers composed of α/β- tubulin heterodimers (Hahn et al., 2019). Immunostaining for β-III Tubulin results in the robust labeling of the axons (Angelin et al., 2024) while immunostaining for myelin basic protein identifies oligodendrocytes myelin sheaths (Yuan et al., 2021).

Monoclonal Anti-β-Tubulin Isotype III antibody (Tub) and polyclonal Myelin Basic Protein antibody (MBP) were used to label axons and oligodendrocytic components in the contralateral intact part of white matter of spinal cord obtained from rats sacrificed in 20 weeks after implantation. The axons in the white matter were organized into parallel sometimes discontinuous arrays and associated with MBP reflecting microstructure of white matter of the spinal cord (Fig. 3A). Ipsilateral rostral part of the spinal cord of the same animal was also populated by the myelinated axons (Fig. 3B) although the regular structure of parallel axons in the white matter was partially disrupted likely as a result of the hemi-excision. A Z-stack transmitted light image of longitudinal section of scaffold having a thickness of 70 μm was obtained by a confocal microscope (Fig. 3Ca). It demonstrates a well-preserved microstructure of tunnels that was damaged along the cutting surface by a slicing procedure resulting in many cracks. A tissue composed of parallel fibers are clearly visible in the flanking regions of the implant suggesting ingrowth of nerves into the microtunnels. Supporting this suggestion, the recipient tissue labeled for Tub and MBP regenerated into the microtunnels of 3D scaffold, readily passing across host/scaffold interfaces (Fig. 3Cb,c). As it was indicated earlier (Fig. 1), the scaffold material reveals a low level of fluorescence when excited at 543 nm. However, at low level of spatial resolution used in Fig. 3Cb, fluorescence of large volume of scaffold material interferes visualization of Tub staining (Fig. 3Cb), while staining for MBP is clearly observed (Fig. 3Cc) since the scaffold material is not excited at 633 nm used for MBP imaging. The latter also allowed to overlay oligodendrocyte staining with one of scaffold material shown in green in Fig. 3Cd. All microtunnels, except ones damaged by slicing procedure, studied in rats (n = 6) in 12 and 20 weeks after the implantation contained the recipient tissue stained for Tub and MBP. The tissue penetrating the scaffold grew in linear orientations along the rostro-caudal axis of the spinal cord, guided by microtunnels; the rostral and caudal endings of the scaffolds were symmetrical. We did not observe a difference in immunostaining of MBP at rostral and caudal recipient/scaffold interfaces of microtunnels (Fig. 3F). This suggests that descending and ascending axons regenerating into microtunnels reach the opposite ends of the lesion site. Unfortunately, it was not possible to obtain transverse cross-sections through implanted scaffolds since the scaffold material was fragile and our attempts to cut the scaffolds using a vibrotome led to their destruction. Thus, the implanted PEGDA/PETA scaffolds retain physical support across a lesion site during 20 weeks after implantations resulting in substantial tissue regeneration into scaffold microtunnels.

**Figure 3.**
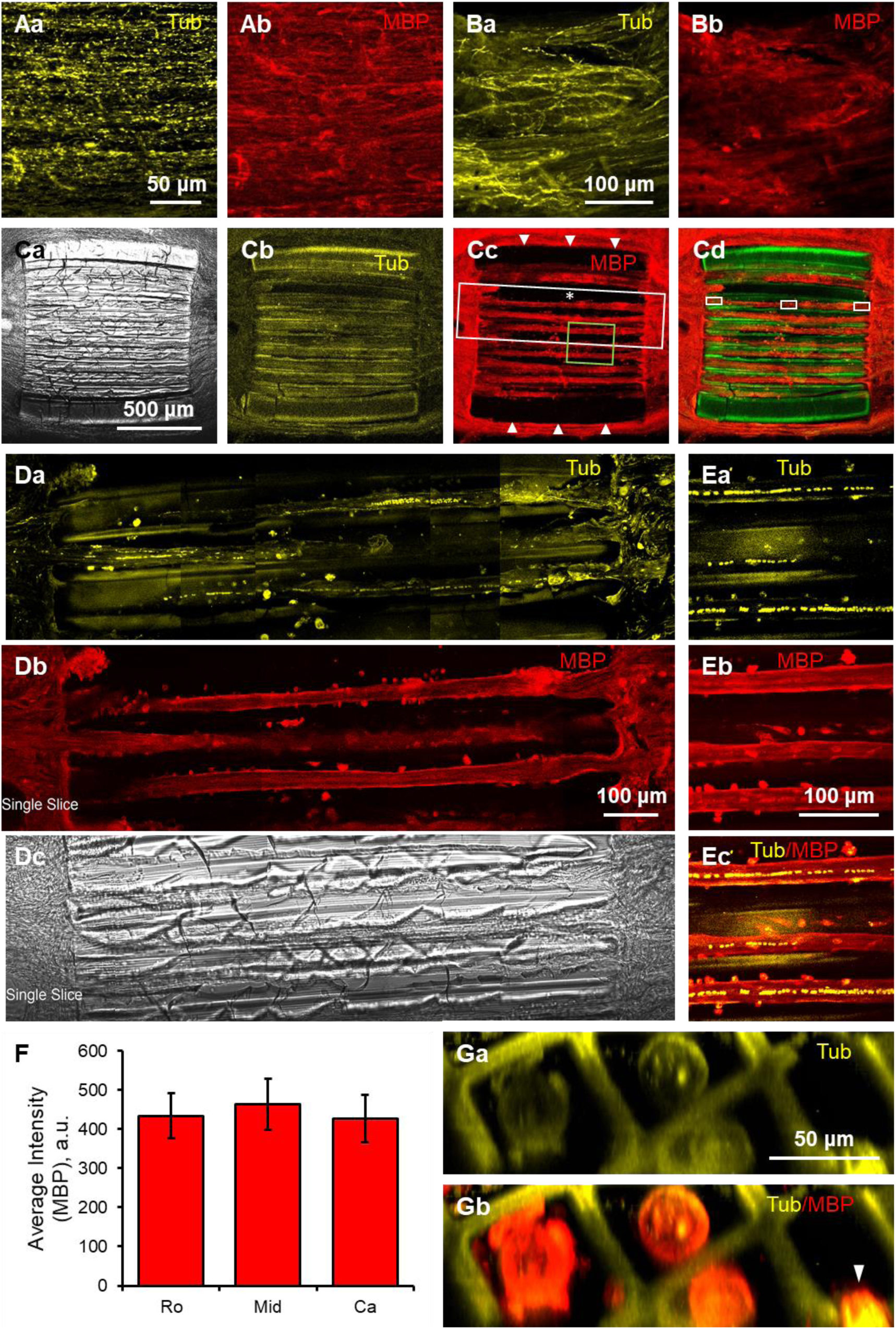
Ingrowth of recipient tissue in 3D-printed microtunneled scaffold in 12 weeks after implantation. **A**. Tub (yellow, **a**) and MBP (red, **b**) labeling of contralateral white matter of T12 rat spinal cord. Rostral is to the left, and caudal is to the right of the image. The axons observed as interrupted yellow lines are organized in parallel and aligned along rostrocaudal axis of the preparation. The MBP staining demonstrates their myelination. **B**. Tub (yellow, **a**) and MBP (red, **b**) labeling of ipsilateral white matter of rat spinal cord rostral to the implantation site. Rostral is to the left, and caudal is to the right of the image. The axons are mainly aligned along rostrocaudal axis of the preparation, although their parallel organization is partially disrupted due to the lesion induced by operation (**a**). The MBP staining demonstrates preserved axon myelination (**b**). **C**. An implanted scaffold harvested in 12 weeks after the implantation. Transmitted light Z-stack image of the slice of scaffold implant (**a**). Note fibers entering the microtunnels from both rostral and caudal ends. A mesh of wrinkles are defects in the scaffold emerged due to slice cutting procedure. Tub (yellow, **b**) and MBP (red, **c**) labeling of the slice of scaffold implant. Yellow fluorescence of stained tissue ingrown in the microtunnels is obscured by the autofluorescence of the scaffold materials while the red fluorescence of stained MBP is clearly visible. Wide black strips at the top and bottom in **c** indicated by white arrowheads show the outer wall of the scaffold. These walls are visible as yellow strips in **b** due to scaffold material autofluorescence. The tissue in one microtunnel indicated by a white asterisk in **c** is lost during the slice cutting. An overlay of MBP staining (red) and scaffold material (green) showing the tissue ingrowth in the microtunnels. Tub (yellow, **b**) and MBP (red, **c**) labeling is also clearly visible at the outer surface of scaffold walls. All images were obtained using x10, N.A. 0.3 objective. White and green boxes are shown with higher magnifications in **D** and **E**, respectively. **D**. Higher magnification and higher resolution Z-stack image (depth of 20 µm) of microtunnel obtained by stitching of Z-stack recorded using x40, N.A. 0.85 objective. Tub (yellow, **a**), MBP (red, **b**) labeling and transmitted light (**c**) images of three microtunnels in the area indicated by a white rectangle in **Cc** are shown. Images demonstrate almost even distribution of stained tissue along the microtunnels. Some interruptions of the tissue along the microtunnels occurs since it was not possible to choose focal planes crossing the tissue along the entire length of all neighboring microtunnels. Substantial damage of scaffold material by slice cutting (**c**). **E**. Z-stack of the area shown in a green rectangle in **Cc** showing even tissue distribution in the microtunnels. Note dotted yellow lines in each tunnel reflecting tubulin staining of erythrocytes shown in more details in Fig. 4E. **F**. A histogram demonstrating similar intensities of MBP fluorescence in first 100 µm at rostral and caudal ends of microtunnels and in its middle part (21 microtunnels from 3 different implanted scaffolds). Examples of regions, from which the fluorescence was quantified, are shown in **Cd** as white rectangles. **G**. Lateral cross-sectional images of several microtunnels obtained from a 3D Z-stack reconstruction. Note the microchannel walls shown in yellow and the protrusion of tissue (indicated by the arrowhead) in one tunnel into the adjacent one due to the splitting of the microchannel wall during sectioning. All images in this figure were taken from the spinal cord and scaffold harvested in 12 weeks after implantation.

Immunostaining of regenerated tissue all along three microtunnels is shown in Fig. 3D with a higher magnification images of the central part of microtunnels depicted in Fig. 3E. The staining is observed all along the microtunnels with some interruptions occurring when the tissue was not in the focal plane of the microscope. Cylindrically shaped MBP labeling is supplemented by staining of cell somas, most probably oligodendrocytes, located on the surface of the cylinders (Fig. 3Db and Fig. 3Eb). Tub immunostaining is observed as rows of large and small dots running all along the microtunnels (Fig. 3Da and Fig. 3Ea) partially overlapped with MBP staining (Fig. 3Ec).

3D reconstruction of sagittal optical section of the implanted scaffold in 100 μm from its end demonstrates robust MBP staining in the central part of the implant forming a cylinder having diameters of about 30 μm (Fig. 3G). The cylinders were attached at their sides to the walls of the microtunnels indicating reasonable adhesion properties of PEGDA/PETA mixture. The rightest microtunnel is damaged by the cutting procedure that results in shifting of the regenerated tissue in the neighboring microtunnel. It explains why sometimes microtunnels seem to be empty as in Fig. 3Cc. The lack of staining for MBP and Tub between MBP-labeled tissue and scaffold walls (Fig. 3G) suggests absence of regenerated nerve tissue in this space. On average, 25–35% of cross-section in the central part of each microtunnel was populated by the regenerated nerve tissue implying a substantial level of possible functional recovery.

Next we employed a high resolution confocal imaging in order to better understand microarchitecture of the stained tissue on the outer surface of the implanted scaffolds and inside their microtunnels. Substantial amount of recipient tissue regenerated over outer surface of scaffolds (Fig. 4A). Both material of scaffold and Tub are visualized in Fig. 4Aa showing their close opposition while MBP staining is overlapped with one of Tub. Higher resolution Z-stack, obtained from the region indicated by a white box shown in Fig. 4Aa, demonstrates a high density of axons growing over the surface of implanted scaffold. Altogether these data prove the properly aligned regeneration of myelinated axons in the superficial layer of at least 50 μm in depth over the scaffold.

**Figure 4.**
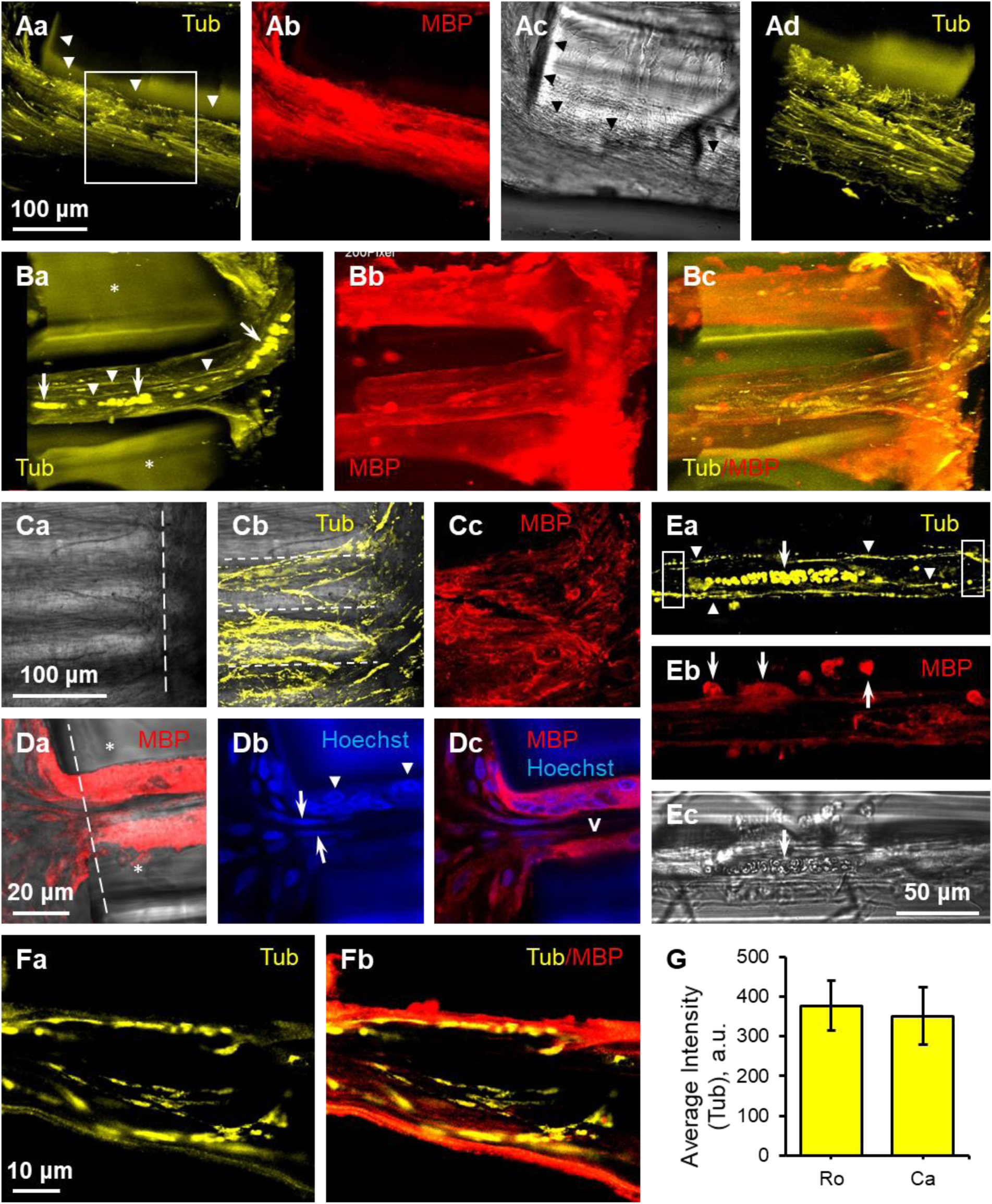
Microtunnels and outer surface of 3D-printed scaffold provide guidance for vessels and myelinated axons. **A**. Myelinated axons regenerating on the outer surface of 3D-printed scaffold. Tub (yellow, **a**), MBP (red, **b**) labeling and transmitted light (**c**) images (Z-stacks) demonstrating regenerated myelinated axons in a layer of about 50–100 µm over the lateral side of the implanted scaffold. The scaffold material is indicated by arrowheads in (**a**) and (**c**). Higher resolution image (Z-stack) of an area indicated by a white rectangle in (**a**) shows numerous axons growing over the outer surface of the scaffold. **B**. 3D reconstruction of the recipient tissue entering scaffold microtunnels. Some axons are indicated by white arrowheads while erythrocytes localized in a vessel are indicated by white arrows (**a**). Scaffold material is indicated by asterisks. MBP immunostaining demonstrating the tissue in three microtunnels (**b**). MBP is visible in three microtunnels rather than in one observed for tubulin staining since the scaffold material is not fluorescent at 633 nm excitation used for MBP imaging. Overlay of Tub and MBP staining is shown in (**c**). **C**. Tissue regenerated in the microtunnels and released when the implanted scaffold was sliced. Transmitted light image of the scaffold located below the focal plane, from which the tissue was released (**a**). A white dashed line indicates the boundary of the scaffold. White dashed lines in **b** outline the microtunnels, from which the axons stained by antiTub antibodies, were released. MBP staining demonstrates that a substantial part of the axons is myelinated (**c**). **D**. A representative example of entrance to the microchannel. An overlay of transmitted light and MBP staining images (**a**). Walls of the microtunnels are indicated by asterisks. A white dashed line indicates the boundary of the scaffold. Hoechst staining reveals nuclei of different shapes likely belonging to oligodendrocytes (white arrowheads) and epithelial cells (white arrows) (**b**, **c**). Unstained area inside the microtunnel indicated by **v** in (**c**) is likely to be a lumen of the vessel. **E**. Axons, erythrocytes, and oligodendrocytes in a scaffold micro tunnel. Erythrocytes (white arrows) aligned along a vessel running close the microtunnel axis and axons (white arrowheads) situated in parallel in the proximity to the capillary stained using Tub antibody (**a**). MBP staining demonstrates somas of oligodendrocytes (white arrows) also situated in a close proximity to the capillary (**b**). A transmitted light image validating that yellow round-shaped spots shown in (**a**) are mainly erythrocytes (**c**). Values of fluorescence attributed to axons were measured in rostral and caudal regions of field of view (shown as white rectangles in (**a**)) for each studied microtunnel. The values for rostral and caudal regions **F**. Z-stack image of myelinated axons in a microtunnel; Tub (**a**) and overlay of Tub and MBP (**b**) staining. **G**. A histogram showing no significant difference in the axonal fluorescence at the rostral and caudal sides of the obtained images (n = 23 images from 3 scaffolds). An example of rostral and caudal regions, in which axonal fluorescence was measured, is shown in **Ea**. All but **E** images in this figure were taken from spinal cords and scaffolds harvested in 20 weeks after implantation. x60, N.A. 1.4 objective was mainly used for imaging.

There was also a strong ingrowth of recipient tissue into the microtunnels of scaffolds (Fig. 4B). 3D reconstructions of first 100 μm from the end of the scaffold reveal several microtunnels with the recipient tissue entering them. Only one microtunnel with Tub staining is visible (Fig. 4Ba) since the tissue in microtunnels located below was covered by their walls, which fluorescence prevents observation of the tissue. Axons entering the microtunnel are clearly visible. Yellow spots of about 5–10 μm aligned over a smooth curve entering the scaffold represent the expression of Tub in cells, most of which are likely to be erythrocytes known to express this protein (Murphy et al., 1986). The smooth curve along which the cells are aligned is associated with a capillary entering the micotunnel. These suggestions will be additionally validated below. Excitation inducing fluorescence of Alexa 647 staining MBP did not excite the scaffold material resulting in possibility to observe MBP staining in two additional microtunnels located below the walls (Fig. 4Bb,c). In many cases scaffolds were substantially damaged by cutting procedure and the tissue was released from the microtunnels laying on the top of scaffold material (Fig. 4C). Many axons entering the microtunnels in the intact implanted scaffold were released from them although still preserving initial bundled microarchitecture. A transmitter light blurry image of the scaffold situated below the focal plane demonstrates microtunnels from which the bundles were released (Fig. 4Ca,b).

MBP-stained recipient tissue grew in linear orientations along the rostral-to-caudal axis of the microtunnel (Fig. 4Da). No cells forming the ‘wall’ at recipient/scaffold interface was observed as evident from staining the nuclei (Fig. 4Db) implying that the used PEGDA/PETA material was biocompatible resulting in an attenuation of reactive cell layers. Importantly, no MBP-stained tissue and nuclei were observed in the central part of the microtunnel (Fig. 4D). At the same time, two different shapes of nuclei, almost round and strongly elongated, were observed at the initial part of the microtunnel. We speculated that the central non-stained part within the microtunnel lacking the tissue and nuclei is a lumen of capillary and the elongated nuclei belongs to cells of the capillary walls. In order to validate this hypothesis, sites within microtunnels revealing strong doted tubulin staining (Fig. 3Ea, 3Fa) were visualized at higher spatial resolution. The staining clearly demonstrates erythrocytes localized to the central part of tissue situated in the microtunnel (Fig. 4Ea). Similar results were obtained for all studied microtunnels having strong doted Tub staining (Fig. 3Da, Ec). It is important to note that the majority of microtunnels in all studied scaffolds had this type of staining in one or several locations along their length. It indicates that a single capillary is likely to pass through almost each microtunnel in 20 weeks after scaffold implantations. The capillaries provide the necessary metabolic support for growth of axons and oligodendrocytes within the microtunnels (Fig. 4E). Immunostaining of extended but discontinuous microtubules using Tub antibodies produced an interrupted pattern for each axon (Fig. 4Ea, Fa) and if parallel axons were bundled together it was almost impossible to trace a single axon throughout the entire length of microtunnel. Additionally, axons were not always passed in a narrow focal plane of used high numerical aperture objectives also resulting in impossibility of long range axonal tracing even in the case if Z-stack imaging was employed. However, no significant difference in the axonal fluorescence was observed at the rostral and caudal sides of the obtained images of the microtunnels as it is shown in Fig. 4Ea, G. Thus, it appears that axons enter the microtunnels of the scaffold from the rostral or caudal aspects of the lesion and regenerate into the opposite side of recipient spinal cord. The axons regenerating in parallel to the capillaries were well myelinated by oligodendrocytic structures (Fig. 4F) situated along microtunnels (Fig. 3Cc, 3Db, 3Eb, 3Gb, 4Da, 4Fb). At least several axons were observed in each microtunnel (Fig. 4Ba, Cb, Ea, Fa). Besides, variability in Tub imunnofluorescence per microtunnel is substantial (Fig. 4G). Even if the lowest level of Tub fluorescence represented only one axon, it also indicated that several axons should populate each microtunnel. Thus, considering that at least 5–10 axons regenerated in each microtunnel and that the scaffold had 169 printed microtunnels, it seems that about 850–1700 axons could potentially regenerate in the scaffold. Suggesting that the axonal density regenerated in the layer of 50 µm of the other surface of the scaffolds (Fig. 4A) is similar to one inside the microtunnels, i.e. about 5–10 axons per each 50 µm length of scaffold circumference equal to 3140 µm, it additionally gives more than 300–600 axons (5–10 axons x (3140 µm /50 µm) = 314–628 axons). Altogether it may lead to at least 1000–2000 axons regenerated throughout the scaffold that creates good opportunities for both motor and sensory partial recovery and reduction of spasticity as a result of scaffold implantation after the SCI.

### Functional recovery after hemi-excision and scaffold implantation

To evaluate whether 3D biomimetic microtunneled scaffolds support functional motor recovery, animals were assessed using the Basso, Beattie and Bresnahan (BBB) locomotor scale (Basso et al., 1995) over a 20-week period after implantation. Animals that received cylindrical scaffolds after hemi-excision revealed significant functional recovery compared to animals of control group underwent hemi-excision only, indicating that the presence of a large internal surface area of microtunnels provides for a spatially oriented substrate for the axonal and vascular growth through the lesion site (Fig. 5A, B). By 12^th^ week, the BBB score reached a median value of 9.0 points ([Q1 = 3], [Q3 = 10.75]) in animals with implanted cylindrical scaffolds, indicating movement of each joint of the hindlimb, in contrast to a median score of 0 points ([Q1 = 0], [Q3 = 3.5]) in animals with the hemi-excision (p < 0.05, Mann–Whitney U test; Fig. 5A). The BBB score was not further changed till the end of experiment (p > 0.05, Wilcoxon matched pairs test) simultaneously being significantly different from one of the hemi-excision group (p < 0.01, polled results for measurements after 8^th^ week; Mann–Whitney U test; Fig. 5A).

**Figure 5.**
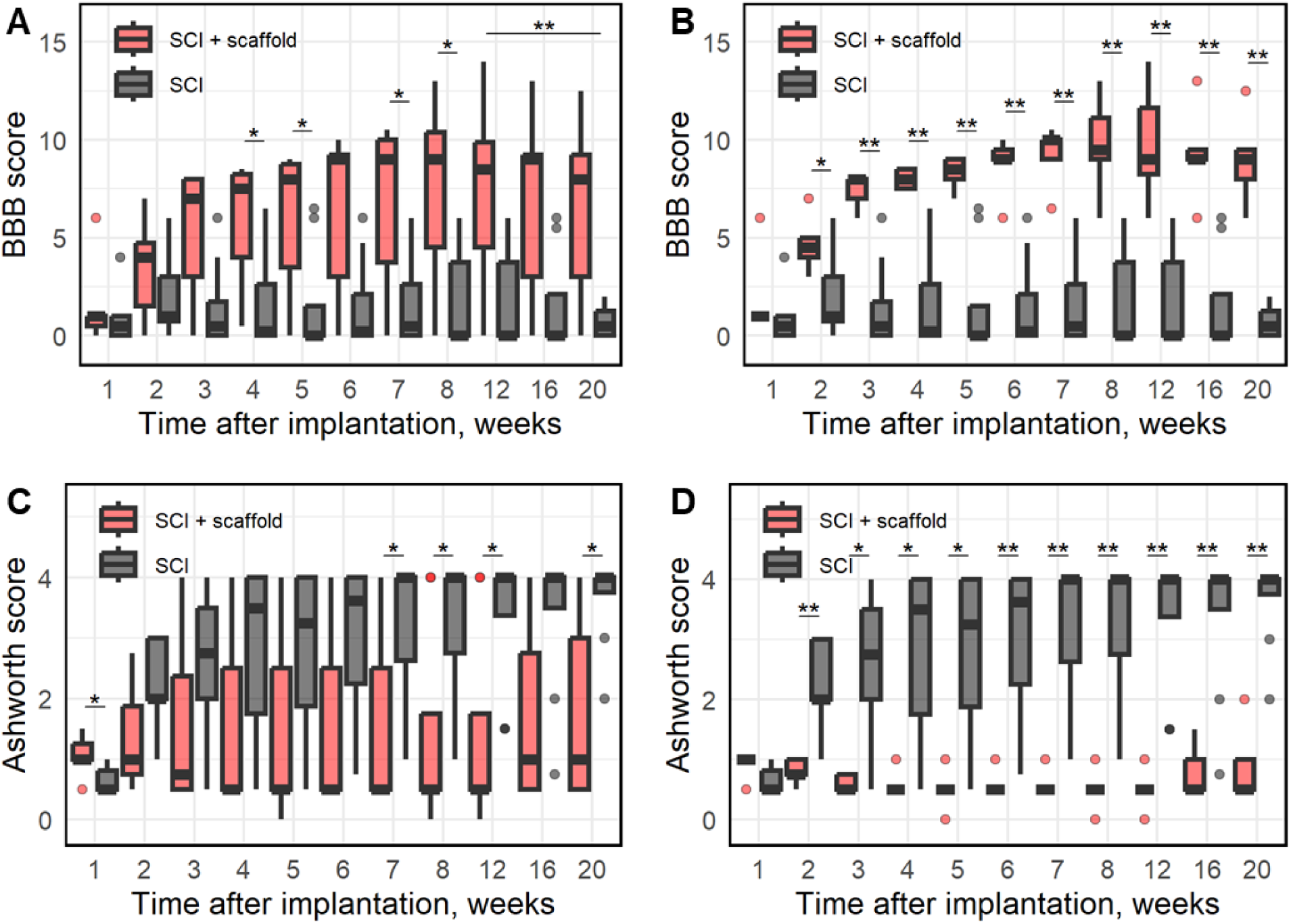
Functional motor recovery and reduction of spasticity after scaffold implantation. **A**. A time course of BBB score changes after the hemi-excision and after the hemi-excision with the scaffold implantation for all tested animals (n = 8 animals in each group). **B**. A time course of BBB score changes after the hemi-excision and after the hemi-excision with the scaffold implantation for all tested animals except two outliers in the scaffold group likely implanted by misprinted scaffolds (n = 6 animals in the scaffold group). **C**. Ashworth scores for assessment of spasticity after hemi-excision and scaffold implantation compared to ones after hemi-excision only. Black and red curves represent Ashworth scores for all tested animals (n = 8 animals in each group). **D**. Ashworth scores for all tested animals except two outliers in the scaffold group likely implanted by misprinted scaffolds (n = 6 animals in the scaffold group). median ± quartile; Mann–Whitney U test; *p < 0.05, **p < 0.01.

In the group with cylindrical scaffolds, in one week after the injury and implantation, the BBB score of the studied hindlimb was 1.4±0.8, being not significantly different from 0.9±0.5 of BBB score for the group with the hemi-excision (p > 0.05, Mann–Whitney U test) indicating that implantation of the scaffolds does not substantially affect the recovery in the most acute SNI period. Then BBB score of implanted animals significantly increased over the next 2 weeks reaching 5.3±1.4 points (p < 0.05 compared to the 1^st^ week, Wilcoxon matched pairs test) further significantly increasing up to 8^th^ week (p < 0.05 compared to the values for both 3rd and 5th weeks, Wilcoxon matched pairs test). The consistent nature of the improvement of the function of the paretic limb in the animals of the scaffold group is also proven by the value of the rank correlation coefficient (r = 0.79) between the averaged values of BBB scores and the duration of observation (р < 0.05, Spearman rank order correlations). It should be noted that one animal of the scaffold group having the highest BBB averaged score at the 8th and 12th weeks (12.75) was sacrificed in 12 weeks to carry out the immunohistochemical research. It led to a drop in the BBB scores in the following observation period (Fig. 5A). However, considering all but this animal, no significant changes in BBB scores were observed in the group after 8^th^ week until the end of the 20-week observation (p > 0.05, Wilcoxon matched pairs test). Thus, our data indicate that the observed recovery of motor function occurred during the first 8 weeks after the scaffold implantation and no deterioration of the function was observed in the following 3 months.

It is important to note that the pooled data of BBB scores (during last three time points of observation) within the cylindrical scaffold group were not normally distributed (p < 0.05, Shapiro–Wilk test). It was due to existence of two outliers having almost zero BBB scores over the whole 20-week period of observation. Most probably the lack of recovery was due some printing defects resulting in microtunnel impassability that resulted in obstruction of axons and vessels regeneration. Printing of 3D scaffolds was accomplished by 200 μm layers from the bottom to the top of the cylinders. Mechanical repositioning of a printer stage in Z-direction could potentially create mismatching between the microtunnels in the neighbouring layers resulting in the impassability of microtunnels. This problem was noticed and sorted out by more precise mechanical manipulations and checking the continuity of the superficial microtunnels as shown in Fig. 1. However, we did not check scaffolds used for the implantation having in mind the problem with their further sterilization necessary for the implantation. Thus, it is potentially possible that a little proportion of scaffolds could have the microtunnel mismatching defects. Therefore, we speculated that we may exclude the outliers demonstrating lack of recovery from the scaffold group. Comparison of functional motor recovery for the control and scaffold groups without these outliers demonstrates the robust improvement of motor function after the SCI induced by the implantation of cylindrical microtunneled scaffolds. The BBB score (9.4 ± 0.7 points) was significantly different from one of the hemi-excision group (1.4 ± 0.4 points; p < 0.001, polled results for measurements after 8^th^ week; Mann–Whitney U test; Fig. 5B).

Thus, implantation of the microtunneled scaffolds significantly increases the value of BBB score during the first two months after implantation and stabilizes the achieved level during the last three months of observation. This means that the scaffolds create conditions for effective tissue repair in the early and intermediate period of SCI ranging from remyelination of nerve fibers to the growth of fibers and vessels through the injury zone (Fig. 3, 4) and possible reinnervation of the caudal populations of motoneurons of the paretic limb.

### Reduced spasticity after scaffold implantation

Spasticity induced by the SCI occur in both animals and humans, resulting in involuntary and sustained contractions of muscles and in pain. It is generally considered that SCI-induced spasticity is due to interruption of descending monoaminergic inputs to the spinal cord finally resulting in an increase in excitability of motoneurons (Elbasiouny et al., 2010). Therefore, we evaluated whether axonal regeneration supported by 3D biomimetic microtunneled scaffolds reduced spasticity induced by the hemi-excision using Ashworth scale in the interpretation of Dong et al. (Dong et al., 2005)(Krotov et al., 2022). One week after injury and implantation in the group with cylindrical scaffolds, the Ashworth score of the examined hindlimb was 1.1±0.1, slightly exceeding the corresponding value of 0.7±0.1 in the group with hemi-excision only. After the first week, during all 5 months, no significant changes in the Ashworth scores were observed in the scaffold group (p > 0.05, Wilcoxon matched pairs test). At the same time, in the control group underwent the hemi-excision, a significant and substantial increase in spasticity to the level of 2.3±0.3 points on the Ashworth scale was noted in the second week of observation (р < 0.05,

Wilcoxon matched pairs test). The entire subsequent period was characterized by a steady statistically significant increase in the Ashworth score ((Fig. 5C; for example, the 7th week relative to the 2nd week, p < 0.05; the 16th and 20th weeks relative to the 5th week, p < 0.05, Wilcoxon matched pairs test). The progressive increase in the level of spasticity of the paretic limb in animals of the control group during the observation was also supported by the value of the rank correlation coefficient (r = 0.99) between the average values of the Ashworth scores and the duration of observation (p < 0.05, Spearman rank order correlations). The final value of Ashworth score in the hemi-excision group was 3.6±0.3 points with 6 out of 8 animals showing the maximum scale value of 4 points. A significant difference between Ashworth scores of the control and scaffold groups was recorded starting from the 7th week of observation till the end of experiment (Fig. 5C, p < 0.05, Mann–Whitney U test). It is worth noting that outliers found in studies of the functional motor recovery (Fig. 5B) were also outliers during spasticity testing. In particular, two animals from the scaffold group demonstrating no motor recovery, likely due to printing defects, showed the highest level of spasticity among the rats of this group (Ashworth score 4.0) by the end of experiment. Excluding these outliers results in the mean Ashworth score for other rats of the group of 0.77 ± 0.12 (mean ± S.E.M.; pooled for all observations after 12 weeks) substantially different from one for the scaffold group of 3.45 ± 0.21 (mean ± S.E.M.; pooled for all observations after 12 weeks; p < 0.001, Student’s t-test; Fig. 5D). Thus, the implantation of cylindrical scaffolds in the site of spinal cord lesion had a solid positive effect on treating spasticity preventing its development during the post-SCI period.

## Discussion

In this study, we focused on how improvements in the physical microstructure of scaffolds enhance spinal cord regeneration and the recovery of motor function in the paretic limb following SCI. For that, we selected a synthetic polymer-based scaffold material (PEGDA/PETA), widely used in the field (Jiu et al., 2024) due to its suitable adhesion, proliferation, diffusion properties, and durability. The scaffold structure was significantly optimized to improve alignment, continuity, porosity, axonal dispersion, and the surface-area-to-volume ratio. Our findings demonstrate that the implantation of these optimized scaffolds at lesion sites led to partial spinal cord regeneration, significant recovery of motor function, and a reduction in spasticity. We concluded that 3D-printed, oriented hollow scaffolds, characterized by a large internal surface area and continuous microtunnels, effectively mimic the natural architecture of spinal cord pathways, creating favorable conditions for axonal regeneration. Moreover, scaffolds with the optimized physical microstructure can be further functionalized with biologically active molecules and supplemented with stem cells (Li et al., 2019)(Rybachuk et al., 2024) to further facilitate the repair following SCI.. These findings provide a foundation for the development of advanced biomimetic scaffolds for regenerative medicine.

### Importance of scaffold microarchitecture for repairing SCI

Ideally, a biomimetic scaffold should reproduce the microarchitected, physical, chemical and biological properties of the native tissue in the SCI site of the spinal cord. In relation to the microarchitecte, it would be preferred for the scaffold to possess small diameter hollow, oriented and continues pathways, occupying the majority of the lesion volume. Thus, the scaffolds should contain microtunnels similar to the size and number of tracts preexisted in the injured site to guide axonal outgrowth. The luminal surface of these microtunnels should be large and smooth as well as to have an adequate adhesion to encourage aligned axonal, cellular and vascular spreading and outgrowth.

In this study, the primary objective in scaffold design was to develop an oriented microarchitecture with minimal material volume and maximal surface area, creating an optimal substrate for axonal and vascular growth. To achieve this, we implemented high resolution 3D printing based on 2- photon microscopy. It allowed us to print walls between the microtunnels as thin as 5 µm significantly decreasing the proportion of material within the volume of scaffold. Excluding the outer layer of the utilized scaffold, the material comprised only 10% of scaffold volume leaving the most part of implanted volume for regeneration of recipient tissue. With employment of implemented materials and developed printing approach, it was the minimal possible thickness of walls. Further reduction of the wall thickness resulted in a change in the square shape of the microchannels and could lead to the wall damage during manipulation of the scaffolds due to the low mechanical strength of the printed product. It seems that in the suggested design we achieved a suitable trade-off between porosity and stiffness to ensure the early stage regeneration while maintaining adequate mechanical properties of scaffolds.

The axons and cells within the matrix require metabolic support. For that, the scaffold material should have optimal diffusional properties, that is especially important at the early stage of tissue regeneration, and the microarchitecture of scaffold should facilitate the fast ingrowth of adequate number of vessels. We earlier demonstrated a high permeability of PEGDA/PETA hydrogel to water-soluble molecules with molecular weights below 1000 (Grebenyuk et al., 2023) that implies the fast diffusion of glucose and other metabolites via the scaffold walls. We have also shown vascularization of each microtunnel within the implanted scaffolds in three months after the operation. Obviously that this vascularization was established at significantly earlier stages of regeneration since it had been completely finished by 12^th^ week after the implantation. We have shown that the nervous tissue inside the implant was located close to blood vessels in order for a diffusion to support the cells and axons at the level similar to one occurring in the spinal cord. Thus, engineering of thin wall microtunnels of 50 µm suitable to transport nutrients, oxygen and metabolic waste throughout the scaffold and promoting fast vessel ingrowth and anastomosis with the host vasculature due to large inner surface area of scaffolds improved viability and density of cells and their processes within the implant.

The maximal density of small sized microtunnels have been for the first time employed in this work. The microtunnels covers 90% of cross-section area of the inner part of the scaffold. The size of the microtunnel cross-section was 50 by 50 µm dividing the implanted region in the small continuous oriented volumes vascularized by a single vessel and creating an adequate metabolic support of regenerating axons and oligodendrocytes. Such small characteristic size of tunnels has not been earlier implemented in the field probably due to a lack of necessary fabrication technologies. Mostly tunnels with characteristic sizes of more than 200 µm were implemented (Koffler et al., 2019) although a smaller size of 125 µm was also used (Vijayavenkataraman et al., 2018). Tunnels in these works were not tightly packed that substantially reduced their cross-section area and as a result decreased the inner surface area necessary for supporting the ingrown tissue. Besides the higher tunnel density used in this study we have also implemented the substantially smaller tunnel size that resulted in a quite significant (many fold) increase in the surface-area-to-volume ratio of the developed scaffolds. For example, scaffolds with tightly packed squared 50 µm microtunnels have the almost 6 times higher surface area compared to a scaffold with tightly packed round 200 µm tunnels (Koffler et al., 2019). Altogether the implemented microarchitecture of the scaffolds resulted increase in the surface area in more than order of value compared to previously employed approaches utilizing scaffolds with continuous oriented tunnels. At the same time, almost hollow design of the scaffold did not prevent the fast recipient tissue ingrowth contrary to scaffolds fabricated using microfibers (Zhao et al., 2023) or different fillers (Singh et al., 2018). An additional and quite important advantage of the design with small-sized continues and oriented microtunnels is a significant decrease in the dispersion of regenerated axons. Our results demonstrate that the axons enter each microtunnel of the scaffolds and are distributed in parallel to a vessel localized in this microtunnel, thus far leaving the scaffold with a diversion not exceeding 50 um. Thus, there is a high probability for regenerating axons of sensory and motor neurons to target the proper ascending and descending pathways at the ends of the scaffold leading to fast and correct reestablishment of the innervation. We consider that morphological results of this work demonstrating the complete vascularization of scaffolds and regeneration of at least several thousand axons throughout the full length of microtunnels strongly support the idea of great importance of scaffold microarchitecture for the successful treatment of SCI.

### In vivo performance of 3D-printed scaffolds

This work demonstrated substantial recovery of motor function following severe SCI. Functional scores reached a mean value of 9.4 ± 0.7 points on the BBB scale in 12 weeks after implantation in animals receiving correctly printed scaffolds. This score indicates movement in all joints of the ipsilateral hindlimb and underscores the critical role of scaffold microarchitecture in facilitating proper nerve tissue regeneration. In contrast, prior work employing 3D-printed PEGDA-based scaffolds with 200 µm round tunnels reported minimal motor recovery (1.6 ± 0.8 points on the BBB scale), (Koffler et al., 2019). Only the addition of neural progenitor cell (NPC) grafts improved outcomes in that study, yielding scores of 6.6 ± 0.5 points. This comparison highlights the importance of functionalizing scaffolds but also emphasizes the significant impact of high-resolution microarchitecture. Although the SCI model used by Koffler et al. (2 mm full excision) differed from the 1 mm hemi-excision in this study, control groups in both works (injuries without scaffolds) showed minimal recovery: 0.3 ± 0.2 points in Koffler et al. vs 1.4 ± 0.4 points (this study). Thus, the decent functional recovery we have shown and its lack with scaffolds having low resolution design is likely attributed to the advanced microarchitecture.

Spasticity is one of the most frequent manifestations of SCI and in the long term the proportion of spinal patients with this disorder in some types of SCI reaches 80%(Holtz et al., 2017). This disorder on the background of SCI is manifested by an involuntary increase in tone and motor reflexes of paretic muscles and their spasms, significantly worsening the quality of life of patients (Biktimirov et al., 2023). The mechanisms of spasticity development in SCI are different, but they all involve disruption of descending regulation of spinal motor neurons due to destruction of long projections of upper motor neurons (Windhorst and Dibaj, 2023). Treatment of spasticity includes limitedly effective conservative and surgical measures that primarily affect denervated segmental neural networks of the spinal cord (Tamburin et al., 2022). Nevertheless, it is obvious that the recovery of the lower motor neuron innervation pathways of the spinal cord, in particular by bioengineering methods, is the most logical means of simultaneously repair motor function and reduce spasticity after SCI (Abolghasemi et al., 2024). Having in mind that about 1000-2000 axons potentially regenerate through the scaffold, it is likely that these axons may innervate the lower motor neurons resulting in prevention of spasticity observed in this study.

Thus, implantation of scaffolds has a tangible positive effect on the course of the recovery process, which is associated with the extension of the recovery process up to the 6–8th week of observation. The duration of this period excludes the connection of the positive effect with the processes of rapid remyelination of demyelinated fibers. In our opinion, it testifies to the realization in a significant volume of another mechanism of regeneration — the medium- and long-range growth of myelinated axons in the injury zone through the microtunnels and outer surface of the scaffolds. The last assumption was confirmed in a series of morphological studies shown in this work.

### Future directions

A mixture of synthetic polymers PEGDA and PETA was used as a scaffold material in this work to fabricate the durable scaffolds necessary for the almost hollow design. However, physical, chemical and biological properties of the material are not optimal and should be improved in the further scaffold development. As a first and simple measure, it seems reasonable to combine synthetic polymers with proteins in the scaffold fabrication (Jiu et al., 2024)(Bedir et al., 2020) (Koffler et al., 2019). Combining collagen and/or gelatin, which resemble ECM and contain bioactive motifs that can actively promote cell adhesion, vessel ingrowth, cell proliferation, and differentiation, with PEG in DLP printing may be a perfect approach to make scaffolds for spinal cord implantation with necessary physical, chemical and biological properties. Being natural materials they are also biodegrade in the body, and due to their ECM-like composition, their breakdown products have minimal side effects (Jiu et al., 2024). Although we did not observe scar formation on the surface and tunnel entrances of PEGDA/PETA scaffolds used in this work, collagen, which reduces the expression of glial fibrillary acidic protein at the injury site could additionally help to avoid glial scars in spinal cord repair after SCI (Jiang et al., 2020). Besides, collagen, gelatin as well as other natural-based scaffolds have poor mechanical characteristics, when printed into a hollow or geometrically complex structure (Sakiyama-Elbert et al., 2012) resembling one used in this work. Furthermore, collagen experiences rapid within a few days in vivo biodegradation in the recipient tissue, which is not enough for long-term spinal cord recovery. Thus, combination of PEGDA/PETA and collagen is likely to be perfect for further development of scaffolds. PEGDA/PETA could be used in high proportion compared to collagen in the outer scaffold layers to provide mechanical properties necessary for convenience of scaffold manipulation during operation and long term stability in the recipient tissue while the higher proportion of collagen in the inner scaffold structures will mimic ECM and will be beneficial for diffusion of molecules and adhesion of axons and vessels.

There is difference between the mechanical properties of white and grey matter (Tran et al., 2023), which can be also mimicked by the different proportions of natural materials and PEGDA/PETA. The spinal cord is composed of white and grey matters consisting of sensory and motor axons and the nerve cell bodies, respectively. Obviously scaffolds used for repairing the SCI should reproduce this differential microarchitecture of the spinal cord providing for recovery of both the axons and neurons. However, this straightforward idea has not been implemented in designs of currently developed scaffolds. It appears that stem cell grafting in sites of SCI is mainly carried out homogeneously over the spinal cord cross section (Rybachuk et al., 2024) or even in tunnels printed for spinal cord tracts (Jiu et al., 2024) rather than being specifically localized to areas where the gray matter was situated before the injury. At the same time, it is well-known that spinal cord neuronal progenitors grafted into the injured adult rat spinal cord self-assemble organotypic, dorsal horn-like domains and that injured adult sensory and corticospinal motor axons retain the ability to regenerate and recognize appropriate targets within such grafts (Dulin et al., 2018)(Kumamaru et al., 2019). All described above imply that fabrication of scaffolds with empty compartments topologically similar to the grey matter to host neuronal progenitor grafts may result in spontaneous restoration of complex spinal cord circuitry after SCI even without a need for additional exogenous guidance. Initial attempts to fabricate three-dimensional scaffolds with cavities (Jiang et al., 2020) and tunnels (Koffler et al., 2019) that simulate the anatomy of spinal cord tracts have been accomplished. However, the respective scaffolds contain scaffold materials in sites where the gray matter should be localized and, therefore, could not host neurons there. Besides, diameters of the cavities and tunnels in the scaffolds were in a range of hundreds of microns not providing a detailed microarchitecture and not having large surface area for fast growth of axons and vessels shown in this work. Although we did not print the separate compartments for the gray matter in the framework of the current research, this task looks quite plausible having in mind the observed scaffold stability and spatial resolution of 2P printing.

The remarkably diverse set of collateral projections from corticospinal neurons to different regions of the spinal cord in rats and rhesus monkeys (Sinopoulou et al., 2022) implies an even more complex corticospinal projectome in humans. These projections terminate at distinct levels of the spinal cord not allowing to use the same scaffolds for repairing SNI occurring in different spinal locations. Besides, many spinal tracts also cross the midline of the spinal cord further complicating its microarchitecture. These tracts are not oriented in the rostro-caudal direction at the midline intersections. Therefore, the simple single axis anisotropic scaffolds used in the current research is hardly suited for reproducing the genuine microarchitecture of the spinal cord. Thus, to create biomimetic scaffolds for a specific injury site and a specific patient, it is necessary to reproduce the exact microarchitecture of the tracts with micrometric accuracy, which currently seems quite possible using further development of 2P and DLP printing technologies.

## Acknowledgements

This research was funded by NRFU grant # 2021.01/0328 to N.V. V.U. is supported by NASU PhD school. The authors are thankful to Mr. Anrdew Dromaretsky (Bogomoletz Institute of Physiology) for the help with preparation of illustration.

## Author Contributions

Conceptualization, S.G., V.M., P.B. and N.V.; 3D printing methodology, S.G. and A.R.; operation & behavior testing, V.M. and I.A.; immunostaining & imaging data analysis, O.R., Y.S., V.U., T.P. and P.B.; validation, A.R., P.B. and N.V.; writing—original draft preparation, Y.S., S.G., V.M., V.U., T.P., P.B. and N.V.; writing— review and editing, Y.S., V.M., A.R., P.B. and N.V.; supervision, A.R., T.P., P.B. and N.V.; project administration, A.R., P.B. and N.V.; funding acquisition, N.V. All authors have read and agreed to the published version of the manuscript.

## Informed Consent Statement

Not applicable.

## Data Availability Statement

The data are available upon reasonable request.

